# Decoding Nucleosome-Depleted Regions: Insights from Epigenetic Marks, Nucleosome Size, and Thermodynamic Modelling

**DOI:** 10.1101/2023.09.20.558658

**Authors:** Kévin Tartour, Jérémy Barbier, Kharerin Hungyo, Fabien Sassolas, Cédric Vaillant, Kiran Padmanabhan, Benjamin Audit

**Affiliations:** Institut de Genomique Fonctionnelle de Lyon, Univ Lyon, CNRS UMR 522442, Ecole Normale Superieure de Lyon, Universite Claude Bernard Lyon 1, 46 allee d’Italie F-69364 Lyon, France; Univ Lyon, ENS de Lyon, CNRS, Laboratoire de Physique, F-69342 Lyon, France

**Author notes:** These authors contributed equally to this work (Kévin Tartour, Jérémy Barbier).

**Keywords:** epigenetic regulation, histone variants, nucleosome-depleted regions, nucleosome organisation modelling, nucleosome wrapping

## Abstract

Elucidating the global and local rules that govern genome-wide nucleosome organisation and chromatin architecture remains a critical challenge. Thermodynamic modelling based on DNA elastic properties predicts the presence of sequence-encoded nucleosome-inhibiting energy barriers (NIEBs) along vertebrate genomes. They delineate *in vivo* nucleosome-depleted regions (NDRs) flanked by 2-3 well positioned nucleosomes. Here, we compared mouse NIEBs to NDRs observed at CTCF binding sites and active TSSs to reveal specific chromatin organizations. We uncover in MNase-seq chromatin profiles the presence of particles of subnucleosomal length specifically positioned at the border of NIEBs with an enrichment of H3.3 and its modification H3.3 S31Ph, whereas the positioning of nucleosomes bearing H3K27ac appears insensitive to NIEBs. Surprisingly, post-translational modifications affect the size distribution of nucleosomes as seen by MNase digestion and so likely their breathing capability. We implemented an extension of our thermodynamic model allowing for variable particle size and suggest that subnucleomes at NIEB borders would result from the recruitment of chromatin remodellers at NIEBs. Our findings provide new insights into the mechanisms by which the DNA sequence and epigenetic marks shape the nucleosome positioning and breathing.

## Introduction

In eukaryotes, genomic DNA is condensed into a nucleoprotein chromatin complex, and its primary unit - the nucleosome - consists of about 147 bp of DNA wrapped around an octamer of histone proteins. On the one hand, by packaging DNA, nucleosomes squeeze the genome within the nuclear space, ensure proper mitotic condensation and segregation, and could in theory act as a driving force for the expansion of eukaryotic genomes. On the other hand, as a natural barrier to all DNA-templated processes, nucleosomes contribute to the emergence and the evolution of new regulatory pathways in transcription, replication, recombination, repair and insertion. Chromatin architecture can further be modulated by DNA modifications, ATP-dependent nucleosome remodelers and the introduction of histone variants. All these factors contribute to stabilize or organize nucleosomal landscapes at several levels - from unwrapping nucleosomal DNA on the linear fiber to regulating nucleosome-nucleosome interactions on higher order chromatin (Lobbia *et al*, 2021).

The role of the genomic sequence in nucleosome formation has been a longstanding question since the discovery of curved DNA (Trifonov, 1985). To understand the intrinsic preferences and contribution of the DNA sequence, *in vitro* experiments using competitive assembly of histones on short fragments have been conducted in the past. The analysis of a large set of natural and synthetic sequences, as well as selection experiments (Selex) (Lowary & Widom, 1998; Widlund *et al*, 1997), found that sequences with the highest affinity for histones exhibit a strong 10 bp periodicity of AA/TT/TA dinucleotide steps. Unlike the well described 601 positioning sequence (Lowary & Widom, 1998), the sequences associated with highly positioned nucleosomes *in vivo* present only a weak 10 bp periodicity and cannot fully account for the well-positioned nucleosomes flanking nucleosome-depleted regions (NDRs) (Chevereau *et al*, 2011).

In genome-wide mapping experiments using *in vitro* reconstituted low-density chromatin in yeast (Kaplan *et al*, 2009; Zhang *et al*, 2009, 2011) and human (Valouev *et al*, 2011) genomes, the main intrinsic determinant of nucleosome formation has been found to be GC content and poly(dA:dT) tracts (Tillo & Hughes, 2009), rather than the 10 bp periodicity. Based on these findings, we have developed a thermodynamic model of nucleosome positioning (Chevereau *et al*, 2011, 2009; Milani *et al*, 2009). This model calculates the sequence-dependent elastic energy cost of wrapping a 147 bp sequence around the histone octamer and incorporates the excluded volume component of nucleosome-nucleosome interactions. The model accurately reproduces the low-density *in vitro* data, validating the sequence-dependent energy model (Chevereau *et al*, 2009; Milani *et al*, 2009). By adjusting the chemical potential to achieve high nucleosome density corresponding to *in vivo* conditions, the model confirms that a significant proportion of observed NDRs in chromatin are associated with sequence-encoded nucleosome-inhibiting energy barriers (NIEBs) from yeast to human (Chevereau *et al*, 2009; Drillon *et al*, 2015, 2016; Brunet *et al*, 2018). However, it is important to note that the correlation between the predicted and observed NDRs is not always exact, indicating that DNA-binding proteins and chromatin modifiers may still have additional roles in shaping native NDRs along with the intrinsic genomic sequence contribution (Zhang *et al*, 2009, 2011).

Interestingly, the thermodynamic model not only reproduces the *in vivo* NDRs at NIEBs but also the periodic pattern of well-positioned nucleosomes flanking them (Chevereau *et al*, 2011; Drillon *et al*, 2015, 2016). This statistical positioning arises as a consequence of confining non-overlapping, stochastically moving nucleosomes next to NIEBs (Kornberg & Stryer, 1988), suggesting a non-local effect is responsible for positioning. The strength of the NIEBs and the nucleosome density influence the ordering effect. *In vivo*, the observed NDRs predicted from the sequence-encoded NIEBs exhibit a weak ordering of 1-3 positioned nucleosomes at their borders, which aligns well with the model’s predictions at high density (Drillon *et al*, 2015, 2016). This is due to the relatively low strength of the NIEBs. In contrast, NDRs resulting from extrinsic processes involving DNA-binding proteins and sequence specific chromatin modifiers, such as those found at constitutively expressed genes (Lee *et al*, 2007), CTCF binding sites (Schwartz *et al*, 2019) or active replication origins (Eaton *et al*, 2010), exhibit stronger depletion and therefore greater periodic ordering.

In our model and more generally, nucleosomes were assumed to have a fixed wrapping size of 147 bp, based on crystallographic studies (Luger *et al*, 1997). However, it is known that nucleosome wrapping length can fluctuate due to the “breathing” of nucleosomal DNA at the entry and exit points, as observed in *in vitro* experiments (Koopmans *et al*, 2009). This breathing can be thermally driven or facilitated by chromatin remodelers, histone modifications and variant histones deposition by histone chaperones (Brahma & Henikoff, 2020; Kono & Ishida, 2020). Unwrapping events can result in partial destabilization, forming subnucleosomal particles like hexasomes, or complete eviction of the octamer. Histone variants and modifications, as well as linker histones, can influence nucleosome dynamics and breathing (Konrad *et al*, 2022; Li *et al*, 2023). The control mechanisms for wrapping/unwrapping dynamics, such as the role of DNA sequence or epigenetic marks, are still largely unknown. Recent advances in paired-end sequencing techniques allow for extracting both the position and size of protected nucleosomal fragments, providing insights into wrapping dynamics. Combining this with MNase digestion at different strengths enables the assessment of nucleosome stability and accessibility throughout the genome and epigenome (Mieczkowski *et al*, 2016; Henikoff *et al*, 2011).

In our study, we extend former studies on yeast and human by analysing nucleosome positioning and wrapping dynamics around NIEBs in mice where more than 1.4 million NIEBs are predicted. These NIEBs correlate with NDRs and induce statistical positioning of 2-3 nucleosomes at their borders, thus potentially establishing nucleosome positions over ∼1/3 of the genome, as previously in human (Drillon *et al*, 2016, 2015). To determine the nature, size, and accessibility of these positioned nucleosomes, we analysed (i) MNase titration experiments where chromatin was digested with several concentrations of MNase, followed by paired-end sequencing, and (ii) MNase digestion of chromatin followed by immunoprecipitation (ChIP) and paired-end sequencing for specific epigenetic signatures. We uncovered the presence of subnucleosome particles strategically positioned at the borders of NIEBs. Surprisingly, these subnucleosomes were not found to be enriched at the borders of NDRs associated with CTCF binding and active TSSs. While no accumulation of histone variant H2A.Z was specifically observed at NIEBs, our findings instead revealed a striking enrichment of histone variant H3.3, at the NIEB borders, which nevertheless required H2A.Z to be expressed in the cell. Moreover, our study demonstrated that the positioning of nucleosomes by NIEBs is contingent upon specific histone tail modifications. While NIEBs acted as barriers for H3K4me1-labeled particles, they failed to influence the positioning of H3K27ac nucleosomes. In addition, subnucleosomes comprising H3.3 S31Ph, exhibited a remarkable overrepresentation overlapping NIEB borders. An extension of our thermodynamic model allowing for variable particle size suggests that the presence of the subnucleosomes at NIEBs border is not directly DNA-encoded but due to chromatin remodelers interacting with NIEBs.

## Results

### Subnucleosome particles are well positioned at the borders of nucleosome-inhibiting energy barriers (NIEBs)

Previously, we developed a physical model that simulated the contribution of DNA sequence to nucleosome positioning, which demonstrated that genomic DNA mainly encodes regions that inhibit nucleosome formation (NIEBs) that then condition the statistical positioning of neighbouring nucleosomes that are parked against the inhibiting regions (Chevereau *et al*, 2011; Vaillant *et al*, 2007). Systematic detection of these regions along the human genome as well as 5 other vertebrate species underlined their ubiquitous presence. The average genome frequency ranges from 1 NIEB every 1.3 kb in Zebrafish (*Danio rerio*) to 1 every 2 kb in Chicken (*Gallus gallus*) (Brunet *et al*, 2018; Drillon *et al*, 2016, 2015). In human, experimental *in vitro* and *in vivo* nucleosome positioning data confirmed that NIEBs correspond to NDRs with 2-3 well positioned nucleosomes at each border (Drillon *et al*, 2016, 2015). These results showed that our physical model captures the histone-DNA interaction with precision and that the intrinsic sequence-encoded nucleosome positioning (NDRs and parked nucleosomes) is pertinent *in vivo* on over ∼38% of the human genome.

In an effort to examine in detail the chromatin organization of NIEB borders, we analysed experimental nucleosome positioning data in the mouse genome at nearly the 1.5M such sites. (1,465,549 NIEB loci). The mean nucleosome density profiles at the 5’ and 3’ NIEB borders predicted by the physical model (Fig 1A and B, green line) are very similar to what was previously observed for human (Drillon *et al*, 2016, 2015), supporting the similarity between the underlying sequence properties on chromatin organization in human and mouse. Examining the corresponding density profiles derived from the deep sequencing of MNase digestion of cross-linked chromatin from mouse embryonic stem cells (mESC MNase 64U dataset, Table 1; see Materials and Methods), we observed that the good correlation with the physical model predictions extends to mouse (Fig 1A and B). NIEBs are NDRs when compared to the surrounding regions and oscillations in nucleosome density profiles is observed at their borders as predicted by statistical positioning. Note that as observed in human, the amplitude of the experimental positioning oscillations is smaller than those predicted by the physical model. The density profiles are identical on either side of NIEBs, thus in all further analyses, we plotted the average density profiles over both sides of NIEBs. The distance to the NIEB borders is counted positively (resp. negatively) for locations outside (resp. inside) the NIEBs.

**Figure 1.**
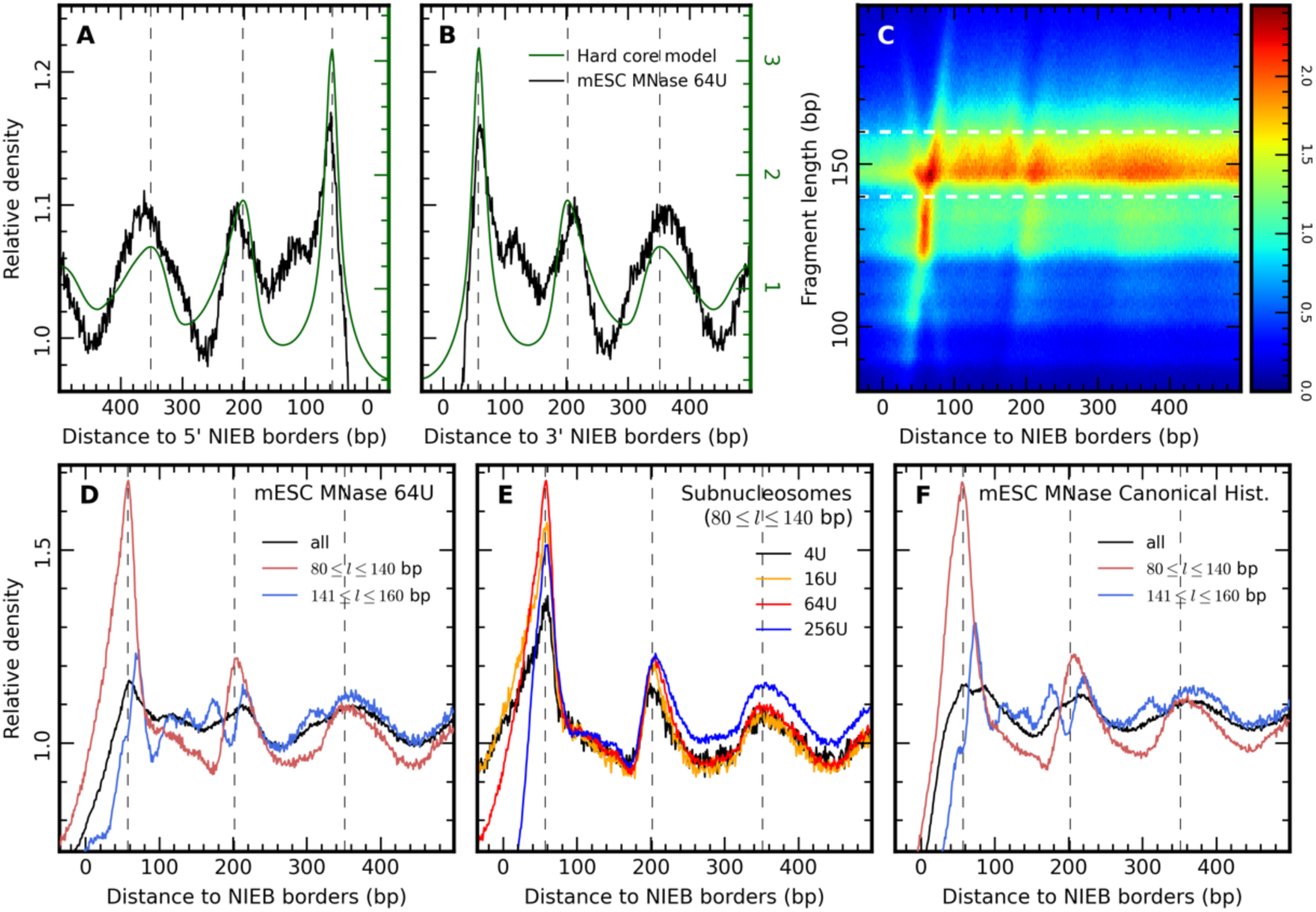
Subnucleosome particles are well positioned at the borders of nucleosome-inhibiting energy barriers (NIEBs) A, B Relative MNase seq density profiles in mESC (mESC MNase 64U, black, Y-scale to the left) and predicted nucleosome occupancy profile (green, Y-scale to the right) at borders of the NIEBs (A, 5’ borders, B, 3’ borders). Predicted nucleosome occupancy is calculated from the energy required to form a nucleosome due to DNA bendability and statistical nucleosome positioning. Both profiles are computed at 1 bp resolution. Negative (resp. positive) distance values correspond to position inside (resp. outside) the NIEBs. C V-plot of mESC MNase 64U data at NIEB borders (2D distribution of MNase fragment’s length and mid-point distance to NIEB borders). X-axis: distance to the NIEB borders, 5’ and 3’ data are merged. Y-axis: fragment length (bp). D Relative MNase seq density profiles at NIEBs borders for all mESC MNase 64U DNA fragments (black), subnucleosome-sized fragments (80-140 bp, red) and nucleosome-sized fragments (140-160 bp, blue). E Relative MNase seq density profiles at NIEBs borders for subnucleosome-sized fragments (80-140 bp) at four concentrations of MNase: 4U (mESC MNase 4U, black), 16U (mESC MNase 16U, orange), 64U (mESC MNase 64U, red), and 256U (mESC MNase 256U, blue). F Relative MNase ChIP seq density profiles at NIEBs borders for canonical histone H3 and H2A (mESC MNase Canonical data) for all, subnucleosome-sized and nucleosome-sized fragments as in D. Note that these density profiles are unchanged when considering mESC MNase H3 and mESC MNase H2A datasets separately (Fig EV1).

**Table 1.**
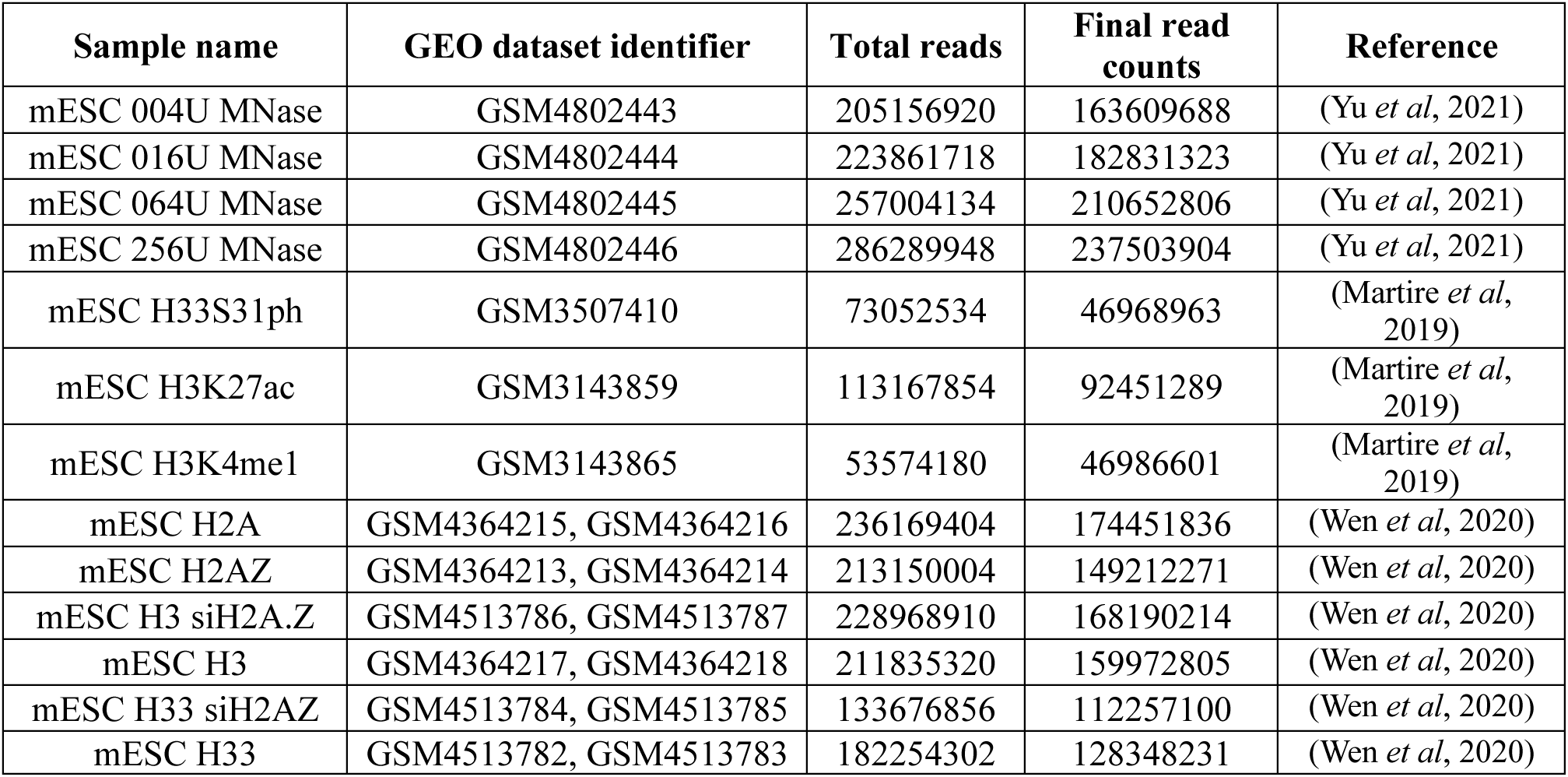
Datasets used in the analyses.

Compared to our previous analysis of human chromatin organization (Drillon *et al*, 2016, 2015), here we selected MNase-seq experiments using paired-end sequencing to interrogate the fragment length of protected DNA at the border of NIEBs (Fig 1C). The V-plot representation (fragment length density plotted against the distance to the NIEBs border) revealed: (i) a peak fragment length around the expected nucleosome length (147 bp) as well as (ii) a population of subnucleosomal fragments specifically at the position of the 1^st^ nucleosome at the NIEBs borders with a length between 100 bp and 140 bp. We further grouped nucleosome fragments between conventional ones (141-160 bp) and subnucleosome fragments (80-140 bp). Their mean density profiles at the NIEBs borders showed that, even though NIEBs are depleted in both particle types, subnucleosome fragments present much stronger preferential positioning compared to full-length fragments (Fig 1D). The density profile of subnucleosome fragments even reproduces the asymmetric wave oscillations predicted by the physical model (Fig 1A and B), especially visible at the 2^d^ and 3^rd^ nucleosome positions (Fig 1D). In conclusion, the physical model predicts the positioning of subnucleosome fragments at NIEBs borders more precisely than full-length fragments.

We then analysed the density profiles for MNase-seq experiments using different concentrations of MNase to digest chromatin (mESC MNase 256U, 16U and 4U; Table 1) and checked whether MNase over-digestion could account for the existence of the subnucleosome fragments (Fig 1E). Independent of the concentration of MNase used (from 16 times less to 4 times more than for the mESC MNase 64U dataset), the subnucleosome fragments presented an identical positioning at NIEB borders. Finally, to test if the DNA fragments we are considering are indeed coming from nucleosomes, we analysed an independent dataset also obtained from cultured mESCs, where MNase digestion of cross-linked chromatin was followed by immunoprecipitation (ChIP) with antibodies specific for either histone H3 or H2A (mESC MNase H3 and mESC MNase H2A; Table 1). The V-plots and fragment length distributions at NIEBs borders (Fig EV2A, D, E and H) as well as the mean nucleosome density profiles at NIEB borders for all DNA fragments, subnucleosome fragments (80-140bp), and full-length fragments (141-160bp) (Fig EV1) were strikingly identical between H3 and H2A profiles. Therefore, we pooled the data together to generate a “canonical nucleosome” dataset (mESC MNase Canonical Hist.). The canonical nucleosome density profiles and V-plot at NIEB borders perfectly reproduced the profiles observed for the mESC MNase 64U data (Fig 1F vs Fig 1D and Fig 2A vs Fig 1C), despite experiments performed in different labs and with or without IP for canonical histones.

**Figure 2.**
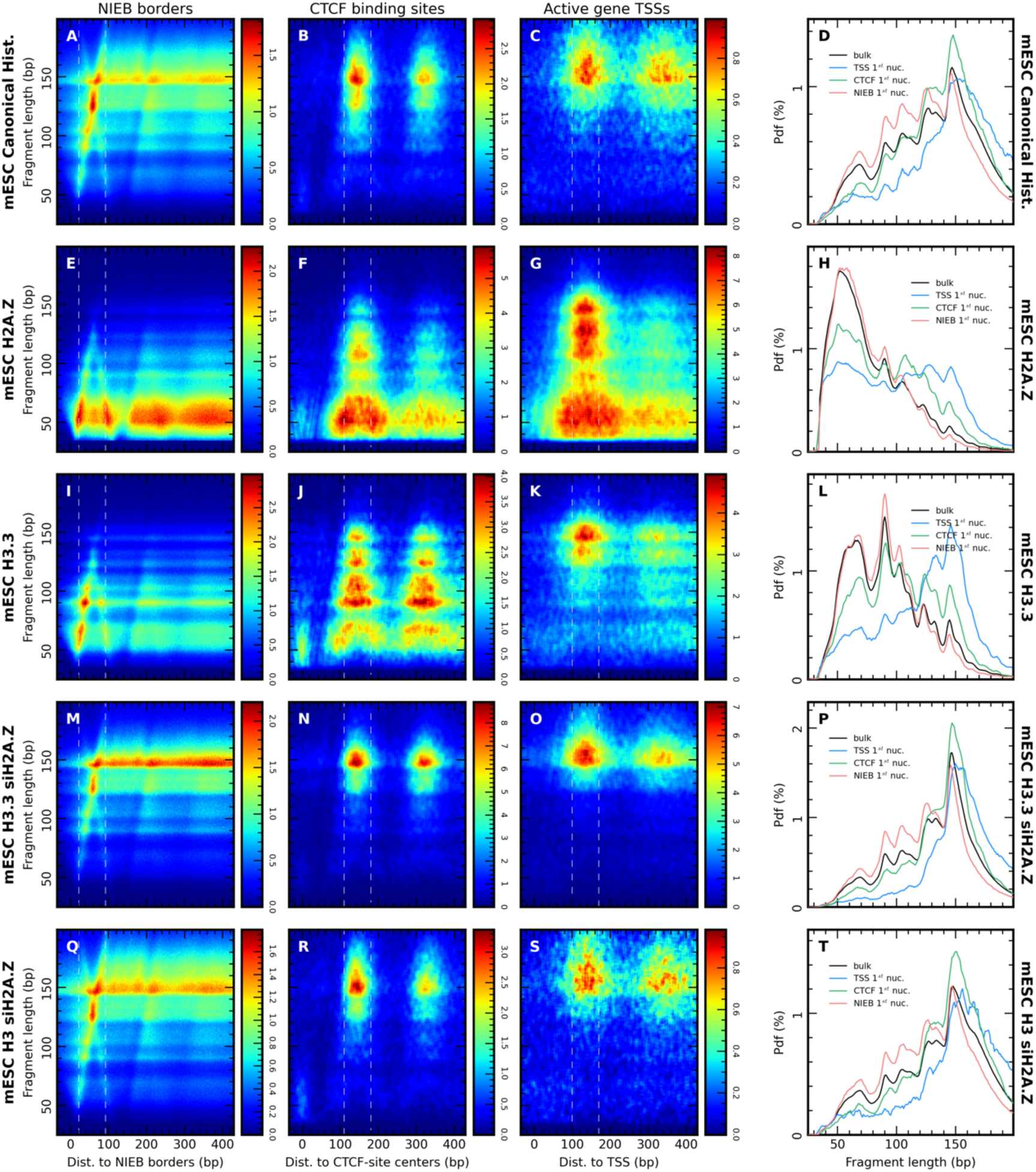
Subnucleosomes at NIEBs borders are independent of histone variant H2A.Z. A-C V-plot of MNase ChIP seq for canonical histone H3 and H2A (mESC MNase Canonical data) at NIEBs border (A), at CTCF binding site (B) and at TSS of active genes (C). X-axis: distance of MNase fragment’s mid-point to the NIEB borders (A), centers of CTCF binding sites (B) and active gene TSS (C). Y-axis: fragment length (bp). In (A) and (B), 5’ and 3’ data are merged. In (C), genes are aligned according to transcription orientation. D Relative fragment length distribution of DNA fragments of mESC MNase Canonical in the bulk (red) or at NIEBs border (red), CTCF binding site (green) and active TSSs (blue). The fragment size in computed at the position of the 1^st^ nucleosome (highlighted with white dotted lines in A-C). Pdf(%): Probability distribution function in percent. E-H Same as A-D for MNase ChIP seq for variant histone H2A.Z (mESC H2A.Z data). I-L Same as A-D for MNase ChIP seq for variant histone H3.3 (mESC H3.3 data). M-P Same as A-D for MNase ChIP seq for variant histone H3.3 in H2A.Z depleted mESC (mESC H3.3 siH2A.Z data). Q-T Same as A-D for MNase ChIP seq for canonical histone H3 in H2A.Z depleted mESC (mESC H3 siH2A.Z data).

Collectively, these analyses provide experimental evidence in mouse that (i) MNase digestion pattern at NIEBs borders can be attributed to nucleosomes/subnucleosomes and (ii) sequence-encoded NIEBs are *in vivo* NDRs associated with 2-3 positioned nucleosomes/subnucleosomes flanking their borders. Furthermore, it revealed that positioning is more salient for subnucleosomes specifically observed at the +1 position immediately adjacent to the NIEB borders. This statistical organization is very robust as observed across datasets from different labs and associated to nucleosome bearing canonical histones.

### Subnucleosome enrichment is not observed at NDR borders associated to CTCF binding and active TSSs

While TSSs and CTCF binding sites are also described as NDRs, they differ from NIEBs in that they are associated with the recruitment of DNA binding proteins such as CTCF and transcription factors. We confirmed the depletion in nucleosomes at these sites compared to the bordering regions when using the mESC MNase 64U datasets but contrary to NIEBs the nucleosome occupancy profiles inferred from the energetic model do not predict this depletion (Fig EV3). NDR at TSSs and CTCF binding sites are note intrinsic nucleosome barriers. Nucleosome depletion is also clearly visible on distribution maps of canonical nucleosome fragment length at these two site categories (Fig 2B and C). Strong nucleosome positioning aligned to these nucleosome barriers was also apparent on these maps with a ∼180 bp nucleosome repeat length (NRL), as opposed to the compact chromatin organization at NIEB borders (NRL<150 bp, Fig 1). Moreover, in contrast to NIEB loci, this positioning mainly involved full-length particles. Comparing probability density function (pdf) of the fragment length computed over the 1^st^ nucleosome position at the NDR borders to the pdf for the full genome (bulk, black line, Fig 2D), it was indeed apparent that the proportion of small particles (<130 bp) is the largest at NIEB borders. Interestingly, particle length distribution at active TSSs is spread to bigger particles compared to the bulk with a larger proportion of particles longer than 150 bp, which could be related to chromatin interactors at these loci. Note that due to robust positioning, the canonical nucleosome density at CTCF borders was very strong, while it was lower than the bulk at active TSSs likely because this nucleosome position is associated with histone variants (Fig 2B, C and Fig EV4A). Hence, the presence of subnucleosomes containing canonical histones at NDR borders is a specific property that distinguishes NIEB NDRs from other NDR sites on the genome, and is not a common feature of all NDRs.

### Subnucleosomes at NIEBs borders are independent of histone variant H2A.Z

The incorporation of histone variants H2A.Z and/or H3.3 into nucleosomes can generate fragile variant nucleosomes which can partially unwrap (Jin *et al*, 2009; Li *et al*, 2023; Wen *et al*, 2020). They are enriched at the +1 and -1 nucleosome positions downstream and upstream of active TSSs and as flanking arrays at CTCF binding sites (Jin *et al*, 2009; Wen *et al*, 2020). We next probed for variant histones in nucleosomes next to NIEB NDRs relative to the other NDRs. The V-plots of H2A.Z-associated MNase fragments at NIEBs, CTCF binding sites, and active TSSs suggested this histone variant is associated with all particle lengths. As previously described (Wen *et al*, 2020): (i) next to active TSSs, where canonical nucleosomes are underrepresented (Fig EV4A), H2A.Z nucleosomes are overrepresented and fragment of all lengths (<160 bp) were observed (Fig 2G, H and Fig EV4B), and (ii) next to CTCF NDRs, over-represented H2A.Z nucleosomes also have a widespread length distribution with fragment length up to 140 bp (Fig 2F and H), full-length fragments being strongly associated to canonical nucleosomes (Fig EV4A). At the 1^st^ nucleosome position next to NIEB NDRs, H2A.Z nucleosomes were mainly observed for particle of lengths <100 bp (Fig 2E and H). These short fragments present a weaker association to canonical nucleosome than for full-length nucleosomes and subnucleosomes (100-140 bp) specific to NIEB +1 nucleosomes (Fig EV4A). The densities of H2A.Z nucleosomes of various sizes at NIEBs +1 nucleosomes are similar to the bulk (Fig EV4B) while their overrepresentation at CTCF and active TSSs +1 nucleosomes decreases with the distance from the NDRs (Fig 2F and G).

Dual variant nucleosomes containing both H2A.Z and H3.3 variants have been associated to nucleosome instability (Jin *et al*, 2009). Accordingly, we observed that the Pearson correlation coefficients between the densities of H2A and H3 nucleosomes and between the densities of H2A.Z and H3.3 nucleosomes are the highest score among the different histone combinations at active TSS and CTCF +1 nucleosomes but also for the bulk and NIEB +1 nucleosomes (data not shown), suggesting that the prevalent configurations at these sites are canonical nucleosomes or dual-variant nucleosome. However, difference could be seen between the H3.3 and H2A.Z fragment length distribution at NIEB loci (Fig2 E and I), H3.3 enrichment at CTCF +2 nucleosomes did not display the decrease observed for H2A.Z (Fig 2F and J) and unlike H2A.Z, H3.3 was not associated to subnucleosomes at active TSS +1 sites (Fig 2G and K). The size distributions of H3.3-associated fragments are skewed towards larger values that H2A.Z-associated fragments. H2A.Z has been described as a major factor associated to subnucleosomes (Li *et al*, 2023). We analysed nucleosome positioning in mESCs where H2A.Z had been knocked down by siRNA (Fig 2M-T, Fig EV4D and E). The decreased amount of H2A.Z abrogates the small size H3.3 particles such that the H3.3 V-plots are close to the original canonical nucleosome V-plots (compare Fig 2M, N, O vs Fig 2A, B, C), with an even sharper distribution around 150 bp of full-length particles (Fig 2D and P, and Fig EV4A and D). The effect of H2A.Z depletion is also slightly visible at the scale of whole H3 nucleosome population, with nucleosome size generally closer from canonical 150 bp nucleosomes (Fig 2Q and T). Apart from particle length distributions, the over/under representations of canonical H3 vs variant H3.3 remains the same (Fig 2M-T, Fig EV4D and E) than in the presence of H2A.Z (Fig 2I-L, Fig EV2E-H, Fig EV4A and C). The observation of the 100-140 bp subnucleosomes associated to NIEB borders is not affected by H2A.Z depletion (Fig 2Q).

Taken together, these results highlight that the implication of histone variants H2A.Z and H3.3 in small-size particles is not identical between the 3 NDR types and the bulk. In this respect, NIEB NDRs were most similar to the bulk chromatin, with an enrichment of histone variants notably at particles with a length <100 bp. Moreover, the 100-140 bp subnucleosomes specifically associated to +1 position at NIEB NDRs are H2A.Z independent as they were still observed for H3- and H3.3-associated particles even upon H2A.Z depletion (Fig 2M and Q).

### Nucleosome positioning by NIEBs depends on histone tail modifications

We next interrogated if certain histone tail modifications could also distinguish the chromatin at NIEB loci. We analysed several MNase profiles in mESC associated to H3 or H3.3 histone modifications (Table 1, Fig 3 and Fig EV5) and found each had a specific particle length distribution and interaction pattern with NDRs. The phosphorylation of Serine 31 on H3.3 (H3.3 S31Ph) that has been linked to heterochromatin formation (Udugama *et al*, 2022; Wong *et al*, 2009) and gene transcription induction (Armache *et al*, 2020; Martire *et al*, 2019) was associated to full-length nucleosomes as well as 110 bp subnucleosomes (Fig 3D). At CTCF binding sites, H3.3 S31Ph nucleosomes were well positioned against the NDR borders and mostly associated to full-length particles (Fig 3B). In contrast, at NIEBs, H3.3 S31Ph nucleosomes were mainly associated to 110 bp subnucleosomes with a preferential positioning overlapping the NDR border (Fig 3A). V-plot for active TSS loci is blurred due to the under-representation of H3.3 S31Ph nucleosomes at these loci (Fig 3C). A strong signal associated to fragment length ∼250 bp was also observed for this MNase ChIP dataset (Fig EV5A-D), possibly resulting from dinucleosomes remaining due to lower MNase digestion efficiency of this chromatin. The analysis of H3K4me1 nucleosomes reveals 3 preferential fragment lengths, ∼110 bp, ∼125 bp and ∼150 bp, whose relative abundance is similar to the bulk across NDR loci (Fig 3H). Moreover, H3K4me1 nucleosomes are well positioned at both NIEB and CTCF NDRs. These nucleosomes were depleted from the active TSS +1 nucleosome position (Fig 3G). The presence of H3K4me1 at the CTCF border and downstream of the TSS +1 nucleosome reproduced previous observations (Barski *et al*, 2007). For H3K27ac nucleosomes, two populations of fragment length are observed in the bulk and at CTCF and NIEB loci: a prevalent population corresponding to full-length nucleosomes (∼150 bp) and ∼130 bp subnucleosomes (Fig 3I, J and L). These two populations are well positioned against CTCF NDRs (Fig 3J). In contrast, H3K27ac nucleosomes appeared not to be affected by NIEB NDRs; they neither presented a strong depletion in these NDRs nor showed positioning at their borders (Fig 3I), reminiscent of the weak positioning of full-length nucleosomes at these loci (Fig 1). At active TSSs, H3K27ac nucleosomes were over-represented and well positioned at the +1 position (Fig 3K). 130 bp and 150 bp nucleosomes were present in equivalent proportion but fragments of length ∼200 bp were also observed with the same prevalence (Fig 3K, L, Fig EV5K and L). To our knowledge, it is the first time that long protected fragments with nucleosomes (200 bp) are revealed associated with H3K27ac at active TSS.

**Figure 3.**
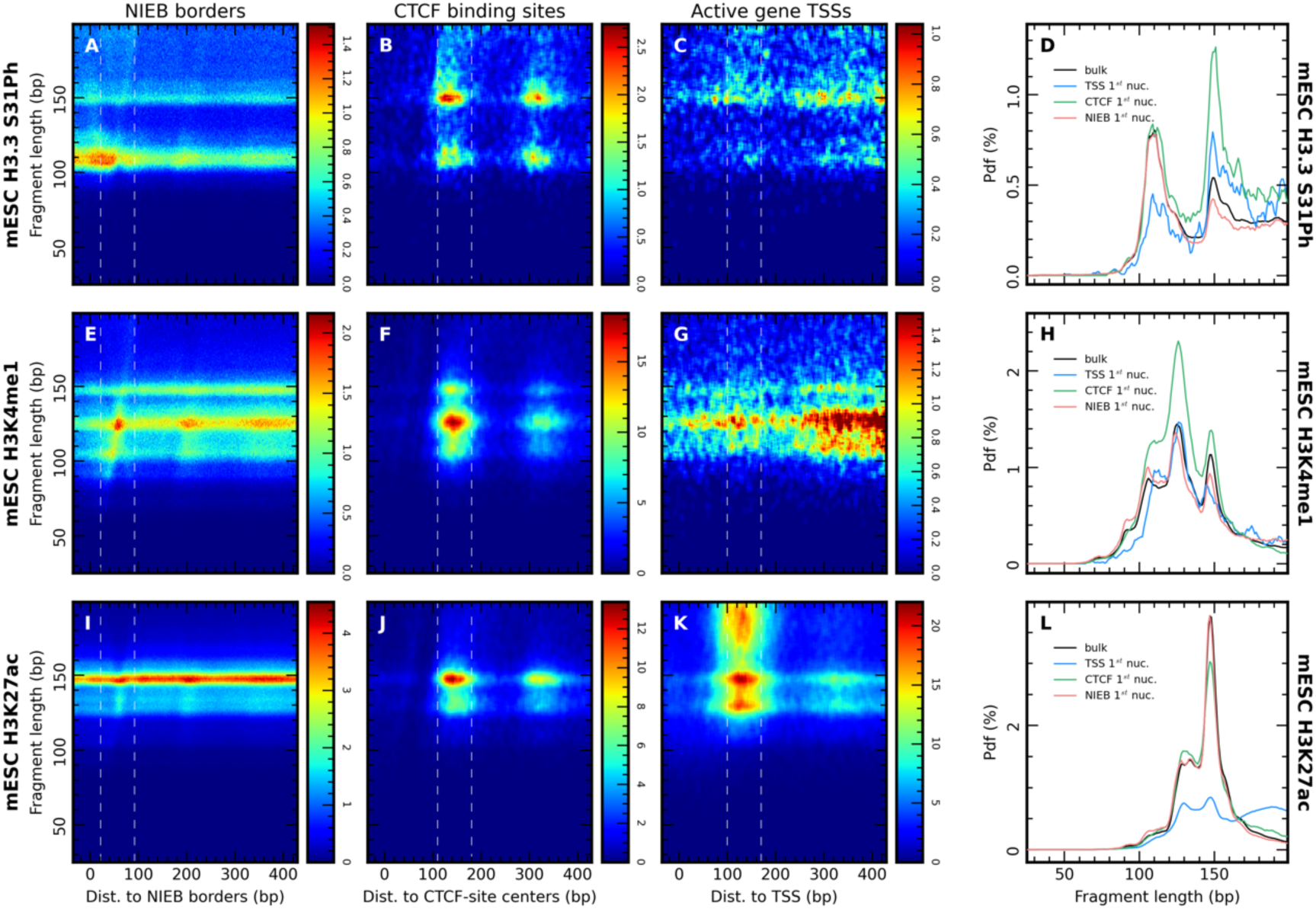
Nucleosome positioning by NIEBs depends on histone tail modifications. A-D Same as Fig 2A-D for MNase ChIP seq for posttraductional modification H3.3 Serine 31 Phosphate (mESC H3.3S31Ph data). E-H Same as Fig 2A-D for MNase ChIP seq for posttraductional modification H3K4me1 (mESC H3K4me1 data). I-L Same as Fig 2A-D for MNase ChIP seq for posttraductional modification H3K27ac (mESC H3K27ac data).

Our analysis revealed that NDRs (TSS and CTCF sites) resulting from DNA-binding proteins were effective positioning barriers for all the analysed particles associated to diverse histone modifications. In addition, they could modulate the size distribution of +1 particles, as exemplified by the 200bp H3K27ac nucleosomes at active TSS borders. Distinctively, the nucleosome positioning effect at sequence-encoded NIEBs depended on the nature of histone modifications. NIEBs acted as positioning barriers for H3K4me1 labelled particles but were ineffective at positioning H3K27ac nucleosomes, while H3.3 S31Ph sub-nucleosomes were over represented at these loci, especially at their border.

### Increased localization of subnucleosomes at the NIEB borders can be explained by remodelling activity

In order to provide an explanation as to how subnucleosomes tend to localise at NIEB NDR borders, we developed a model of nucleosome positioning at thermodynamic equilibrium allowing nucleosome breathing (wrapped DNA length ranging from 91bp to 147bp; see Materials and Methods). The DNA was modelled without any sequence specificity and we fixed the parameters so that nucleosome coverage in the bulk was 80% and that the nucleosome with the longer wrapped length were the more probable (Fig 4B) as observed *in vivo* (Fig 1C). When adding a single 200 bp NIEB in the middle of the bulk DNA corresponding to a 5 kT energy barrier for full length nucleosomes, we observed as expected that NIEBs could position the nucleosomes in regular arrays (Fig 4A). The fragment-size distribution of the +1 nucleosomes at the NIEB borders overlapped almost exactly the bulk distribution (Fig 4B). Changing the height of the NIEB from 5 kT to 2 kT or 10 kT only decreased or increased the positioning but did not modify the fragment-size distribution of the +1 nucleosome that remained identical to the bulk (Fig EV7A and B, and Fig EV7C and D, resp.) This indicated that NIEBs alone were not sufficient to explain the specific presence of subnucleosomes flanking their borders (Fig 1C). Reasoning that the NDRs at the NIEBs could favour the presence of remodelling factors, we then incorporated in the model a size-dependent remodelling activity that destabilized more efficiently +1 nucleosomes when they are fully wrapped. We found that this adjustment was enough to observe a larger proportion of subnucleosomes (Fig 4C) as reflected in the size distributions (Fig 4D). Increasing the strength of the remodelling activity only increased the frequency of subnucleosomes (Fig 4E, F) at the +1 position. Thus, similar to what was experimentally demonstrated, a model that incorporates the +1 nucleosomes remodelling is able to reproduce the excess of subnucleosomes at the NIEB borders compared to the bulk.

**Figure 4.**
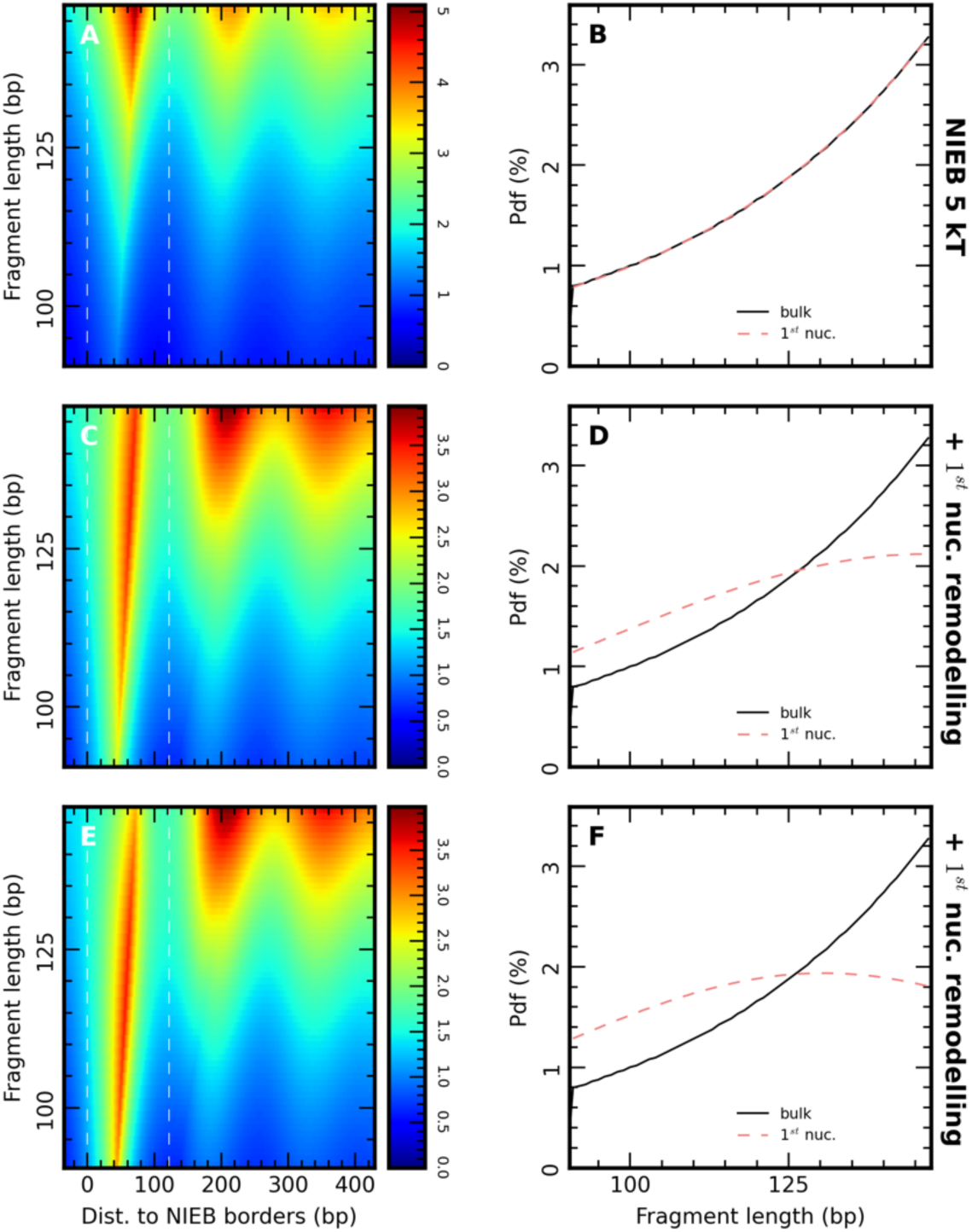
Subnucleosome particles at NIEBs borders could result from specific remodeling activity. A Computed nucleosome dyad-density *P_l_*(*i*) equivalent to the V-plots computed for the MNase data (Fig 1C) at the border of a NIEB of height ℎ = 5 kT positioned within a region with no sequence specificity. The model parameters are set so that nucleosome occupancy in the bulk is 80% (∈_0_ = 0.057 kT/bp and µ = µ*_bulk_* = −10.6 kT). X-axis: distance to the NIEB 3’ border. Y-axis: fragment length (bp) B Fragment size distribution for the model shown in A in the bulk (black) and over the +1-nucleosome position at the NIEB border (highlighted with white dotted lines in A). C, D Same as in (A, B) but with additional specific remodeling of strength: = 1.12 kT. E, F Same as in (A, B) but with additional specific remodeling of strength: = 2.24 kT.

## Discussion

The organization of nucleosomes within the chromatin fibre plays a critical role in DNA compaction. Traditionally, chromatin studies have employed a functional-driven approach, focusing on transcribed regions or transcription factor binding sites to investigate chromatin dynamics. Previous models have highlighted the significance of DNA sequences in nucleosome organization (Awazu, 2017; Chevereau *et al*, 2011; Tompitak *et al*, 2017a; van der Heijden *et al*, 2012). In our study, we adopted a bottom-up approach based on intrinsic DNA sequence-encoded characteristic of chromatin, to uncover novel functions of the genome. We focussed on the 1.4 million NIEBs identified in the mouse genome based on intrinsic nucleosome density profiles predicted by our thermodynamic model that relies on sequence-dependent DNA bendability (Brunet *et al*, 2018). We observed that at NIEB borders predicted nucleosome density profiles aligned with experimental profiles derived from MNase data in mESC. We, thus, confirmed the model accuracy in mice, similarly to previous comparison in human, yeast and other species (Chevereau *et al*, 2009, 2011; Drillon *et al*, 2015, 2016). Published MNase datasets using paired-end sequencing allowed us to deploy the experimental nucleosome density profiles according to MNase fragment size. We discovered an enrichment of subnucleosomal particles specifically at NIEB borders, which remained consistent across different experimental datasets from different labs (Wen *et al*, 2020; Yu *et al*, 2021). Positioning at the border of the NIEB was significantly more pronounced for these subparticles than for full length nucleosomes. Investigating the involvement of histone variants H2A.Z and H3.3 in particle of various size, we confirmed that both H3.3 and H2A.Z are mainly associated to the smallest particles (< 100 bp) and found that small H3.3 particles are overrepresented at NIEB borders. Additionally, we observed that nucleosomes associated to different post-translational histone modifications (PTMs) could escape the general positioning pattern at NIEB borders. Our findings support the relevance of NIEBs, which are intrinsic nucleosome-depleted regions found across species and under selective pressure (Barbier *et al*, 2021; Drillon *et al*, 2016; Brunet *et al*, 2018). The nature of nucleosomes and their short NRL around NIEBs, combined with the DNA-encoded nucleosome depletion, suggests functional importance and active regulation for these regions.

In this context, we know that NIEBs are enriched in poly(dA:dT) tracks (Brunet *et al*, 2018). The presence of these specific sequences may help nucleosome eviction, by enhancing « DNA breathing » at the entry/exit of the nucleosome and promoting the primary fixation of DNA binding proteins (Polach & Widom, 1995) as well as by remodelling factors (Kubik *et al*, 2018). Such DNA breathing was not accounted in our previous equilibrium nucleosome model where the length of DNA wrapped in a nucleosome was fixed (147 bp) (Chevereau *et al*, 2011). In particular, in our previous model, an unfavourable sequence or the presence of a strongly bound complex (a fixed barrier) resist nucleosome formation and cannot induce a partial unwrapping, even though the latter is an appealing scenario to explain the very compact positioning of the 2-3 nucleosomes flanking NIEBs (NRL < 150 bp). Given that the assays performed at different MNase digestion levels indicate a complex picture with NDRs being actually flanked by positioned subnucleosomes, a likely scenario is that the sequence at NIEBs enhances spontaneous nucleosomal DNA breathing leading to more susceptible nucleosomes. Accordingly, it has been recently proposed that such sequence-dependent nucleosome breathing and ATP-dependent remodelling are likely to play a key role in the control of accessibility to buried sites by DNA unpeeling (Isaac *et al*, 2016), and in fine in the control of nucleosome stability to various perturbations. In that respect, our study suggested that accessibility of NIEBs to active chromatin remodelers might play a role in shaping subnucleosomal particles at NIEB borders. Incorporating nucleosome breathing (from 91bp to 147bp) and a particle-size-dependent remodelling activity into an extension of our model successfully reproduced the specific presence of subnucleosomes at the NDR borders, indicating that NIEBs could be preferred sites for chromatin remodelling.

Parallel analyses at alternative *in vivo* NDR loci, active TSS and CTCF binding sites, revealed that NIEBs’ chromatin organization is specific to these intrinsic NDRs, probably due to the mechanism of nucleosome depletion that does not result from competition of histones with other DNA binding proteins, thus favouring the access to other actors such as remodelers. H2A.Z and H3.3 were found flanking the NDRs; but, between NDRs categories (NIEBs, active TSS and CTCF binding sites), nucleosomes exhibited different characteristics, particularly in terms of the length of DNA protected by the nucleosome. In contrast to NIEB loci, nucleosome associated to histone tails modifications were positioned at the preferential nucleosome positions when present. This indicates that nucleosome depletion is not the only player in the specific chromatin organization at NDRs and additional mechanisms must participate. On the contrary, the nature of the nucleosome depletion would contribute to the specification of the neighboring chromatin organization. Chromatin remodelers, such as CHDs, P400, INO80, and SWI/SNF complexes, have been implicated in NDRs formation and organization at TSS and CTCF binding sites (Lorch *et al*, 2014; Clarkson *et al*, 2019). Our study suggests related mechanisms may be at play in the chromatin organization of NIEBs, although further investigation is needed to fully understand these modalities. The RSC chromatin remodeler is known to act on poly(dA:dT) tracts in *Saccharomyces cerevisiae* (Krietenstein *et al*, 2016; Lorch *et al*, 2014). Due to poly(dA:dT) enrichment at NIEB borders (Brunet *et al*, 2018), its mammalian homologous, BAF complexes, could be a good candidate to play a role in the nucleosome depletion and the nucleosome/subnucleosome organization at the borders (He *et al*, 2020). Based on the structure of nucleosome-bound human BAF complex, two models of remodelling are possible. In the first one, a strong DNA translocation induces DNA tension and partial unwrapping leading to nucleosome eviction. In the second model, two nucleosomes enter in collision due to the translocation leading to nucleosome eviction. This could explain that at NIEBs borders the first nucleosome is partially unwrapped, the NRL is particularly short (< 150 bp) and full length nucleosomes are only weakly positioned. Additionally, when one nucleosome has been evicted, cell sensors detects the free DNA and induce its chromatinisation. Histone chaperones could contribute to filling the gaps of free DNA in NIEBs / at NIEB border by depositing available histones. As this process can happen outside of the DNA synthesis period, the available histones are DNA-synthesis-independent histone-variants like H3.3 or H2A.Z but other could be involved. In the case of H2A.Z, its incorporation could also participate to explain subnucleosome particles at NIEB borders as H2A.Z has been shown to modulates nucleosome dynamics by increasing unwrapping (Li *et al*, 2023). Those hypotheses suggest that NIEBs may be the place of high nucleosome turnover.

From an evolutionary point of view, NIEBs are sequence-encoded NDRs that are ubiquitous to all vertebrates with similar characteristic such as their sizes, inter-NIEB distances or GC content oscillations in phase with nucleosome positioning at NIEB borders (Brunet *et al*, 2018; Drillon *et al*, 2016). Strong signs of selection reinforcing the GC content oscillations at NIEB borders have been observed in the human lineage (Drillon *et al*, 2016). These findings strongly suggest a functional importance for these regions. Sequence-encoded NDRs have already been observed having such importance, and even being major driver of speciation (Barbier *et al*, 2021). In yeast, the presence/absence of intrinsic NDRs at the promoter of growth and stress genes differs according to the ideal metabolic pathway of each specie (Field *et al*, 2009). Aerobic yeast exhibit intrinsic NDRs at the promoter of genes associated to respiration, whereas anaerobic yeast exhibit intrinsic NDRs at the promoter of genes associated to fermentation (Field *et al*, 2009; Tsankov *et al*, 2010). The prevalence of intrinsic NDRs at the promoter of genes has also been linked to organism complexity (defined here by the estimation of the number of different cell types in an organism), with the more complex organism having « closed » promoters with nucleosome attracting sequences and simpler organisms having « open » promoters with sequence-encoded nucleosomal barriers (Tompitak *et al*, 2017b; Arneodo *et al*, 2018). These findings highlight the importance of intrinsic NDRs during evolution. Here, we discussed the association of NIEBs with several histone variants and histone PTMs, trying to decipher the functional effect of these sequence-encoded NDRs. As NIEBs have been observed across vertebrates, a clade with a wide variety of species, these functional effects could differ from one specie to another.

NIEBs are homogeneously distributed throughout the genome but display constitutive heterogeneity in their location, barrier length, and strength (Brunet *et al*, 2018; Drillon *et al*, 2016, 2015). By further characterizing NIEBs and incorporating multiple parameters such as nucleosome size at the border, presence of histone variants, and PTMs, it may become possible to categorize NIEBs into homogeneous groups to facilitate the inference of their functional roles among one species or across species. These accessible regions of chromatin could have important implications for essential genome functions such as DNA replication, DNA repair, and 3D structural organization. Future studies will shed more light on the precise functions and mechanisms associated with NIEBs and their impact on genome-wide chromatin organization.

## Materials and methods

### Datasets

For MNase titration data, we used the reads provided by Yu et al. (Yu *et al*, 2021). MNase-ChIP datasets were taken from Martire et al. (Martire *et al*, 2019) and H2A.Z / H3.3 datasets from Wen et al. (Wen *et al*, 2020). All these datasets are paired-end sequenced. Raw reads were downloaded from the SRA bank using the SRA toolkit. GEO dataset identifier are provided in Table 1.

### Pre-processing and alignment

For all the datasets, quality check of sequencing was obtained using the FastQC tool provided by Babraham Institute (Andrew, 2010). FastQC detected the presence of adapters in some of the datasets, that were removed with cutadapt (Martin, 2011) with the following parameters: «-e 0.1 -n 2 -m 35 -q 30 --pair-filter=any ». Adapter sequences used were either found in Illumina documentation based on FastQC detection, or directly given by FastQC. After cleaning, reads were aligned with bowtie2 (Langmead & Salzberg, 2012) with default parameters. Duplicate reads are marked and removed thanks to Picard MarkDuplicates. Only properly-paired reads with a MAPQ of at least 20 were retained for further analysis. The datasets from Wen et al. contain duplicate experiments that are pooled for analysis. Final read counts are in Table 1.

### Post-processing

The position of each fragment was extracted from the alignment file using in-house scripts. This allowed to 1) assign to each fragment a « reference position » as its center and 2) access the length of each fragment.

### V-plots

Length and positions of fragments were compared to the positions of NIEBs, center of CTCF binding sites and TSS of active genes using python3 scripts to determine the V-plots in figures 1-3, EV2 and EV5. We computed the number of fragment of each length falling at each position relative to the NIEBs. The counts were divided by the number of NIEBs considered for each relative position. Finally, this number was normalized by the average density of fragment on the genome, computed as the number of fragments used divided by the total length of the sequenced genome.

### Nucleosome density profiles

Nucleosomes density profiles in figures 1A, B, D, E, F and EV1 were obtained by computing the relative position of each fragment centre to the closest NIEB. At each position relative to NIEB, we counted the number of fragments falling at this position, and divided by the number of NIEBs considered. This number was then normalized by the average read density on the genome, computed as the number of fragments used divided by the total length of the sequenced genome.

### Detection of Nucleosome-Inhibiting Energy Barriers (NIEBs) from the genomic sequence

We use a simple physical model of nucleosome assembly that was shown to mimic *in vitro* nucleosome occupancy data remarkably well in several organisms, including human, yeast and *Caenorhabditis elegans* (Brunet *et al*, 2018; Chevereau *et al*, 2011, 2009; Drillon *et al*, 2016; Milani *et al*, 2009; Vaillant *et al*, 2010). The model consists in computing the sequence-derived free energy cost of bending a DNA fragment of a given 147 bp sequence from its natural curvature to the final super-helical structure around the histone core. Given the resulting energy profile and an average nucleosome coverage (chemical potential), we can derive the nucleosome occupancy profile assuming nucleosome behave as 147 bp hard-core particles. Combining the nucleosome occupancy probability profile and the original energy profile, NIEBs are identified as the genomic energy barriers that are high enough to induce a NDR in the nucleosome occupancy profile as was initially experienced for the yeast genome (Milani *et al*, 2009). When applied to the mouse genome, this methodology predicted 1,465,549 NIEBs having the same genomic properties as those obtained across 6 vertebrate genomes (Brunet *et al*, 2018).

### Modelling partially wrapped nucleosomes at NIEB borders

In order to take into account that some nucleosomes are not fully wrapped, we developed a model where nucleosomes are hard-core particles of variable sizes given by the wrapping length *l* ∈ [*l*min=91 bp, *l*max=147 bp] and can bind to a genome of length L with an additive energy model. The energy to assemble a nucleosome of wrapped length *l* between positions *i* and *i+l-1* is 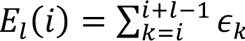, where ∈*_k_* is is the energy cost to include base-pair *k* in a nucleosome. Given the matrix of *E_l_*(*i*) and a chemical potential µ, the equilibrium nucleosome distribution matrix) *P_l_*(*i*) describing the probability to observe a nucleosome of size *l* at position *i* can be exactly computed following the works of Chereji and Morozov (Chereji & Morozov, 2014). Setting all ∈*_k_* to the same value −∈_0_ results in a *bulk* model without any sequence preference. We fixed ∈_0_ = 0.057 kT/bp so that particle wrapping length distribution increases with *l* as observed experimentally (Fig 2D and Fig EV7B, D). Then, the chemical potential was set to µ*_bulk_* = −10.6 kT, so that the mean nucleosomal occupancy be 80% as observed experimentally. The mean density of this bulk model is 1 particle per 157 bp. In this framework, a NIEB could be easily modelled as a gate-function of width 200 bp in the base-pair energy cost profile (Fig EV6A): along a NIEB, the energy cost is 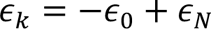 where 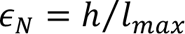 with h the energy barrier height for a full nucleosome. The energy matrix for sequence with one NIEB and the resulting nucleosome distribution map for µ = µ*_bulk_* are illustrated Fig EV6B and Fig 6A, C, respectively.

### Size-dependent remodelling activity

We assumed that the +1 nucleosomes flanking the NIEBs are subject to a remodelling activity preferentially affecting full length particle. To take into account this assumption, the energy cost profile over the 147 bp regions flanking the NIEBs is subject to an additional remodelling term dependent on 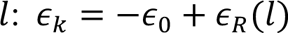, where 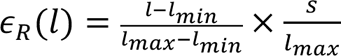 with s the maximal remodelling strength for full length nucleosomes.

## Acknowledgments

We thank lab members and colleagues at the IGFL for their valuable feedback. We thank Alexandre Morozov and Ralf Blossey for inspiring discussions. We thank Zengqi Wen and Guohong Li for sharing their data. This work was funded by the Agence Nationale de la Recherche (ANR-20-CE12-013) to BA and KP. BA and CV acknowledge financial support from the CNRS through the MITI interdisciplinary programs (Miti AAP PIB 2023). The authors thank the French CNRS network GDR “Architecture et Dynamique du Noyau et des Génomes” (ADN&G) for stimulating workshops.

## Author Contributions

K.P., B.A. and C.V. conceived the project with inputs from K.T. and J.B. B.A., C.V., K.P. and K.T. directed the project. K.T., J.B. and F. S. performed the analyses. H.K. developed the physical model with the inputs of B.A. and C. V. K.T. wrote the manuscript with inputs from K.P., B.A. J.B. and C.V.

## Data Availability

The datasets used in this study are available in the following databases:

MNase-seq data: Gene Expression Omnibus GSE158545 (https://www.ncbi.nlm.nih.gov/geo/query/acc.cgi?acc=GSE158545)

ChIP-Seq data: Gene Expression Omnibus GSE146082 (https://www.ncbi.nlm.nih.gov/geo/query/acc.cgi?acc=GSE146082)

ChIP-Seq data: Gene Expression Omnibus GSE114548 (https://www.ncbi.nlm.nih.gov/geo/query/acc.cgi?acc=GSE114548)

---

**Figure EV1.**
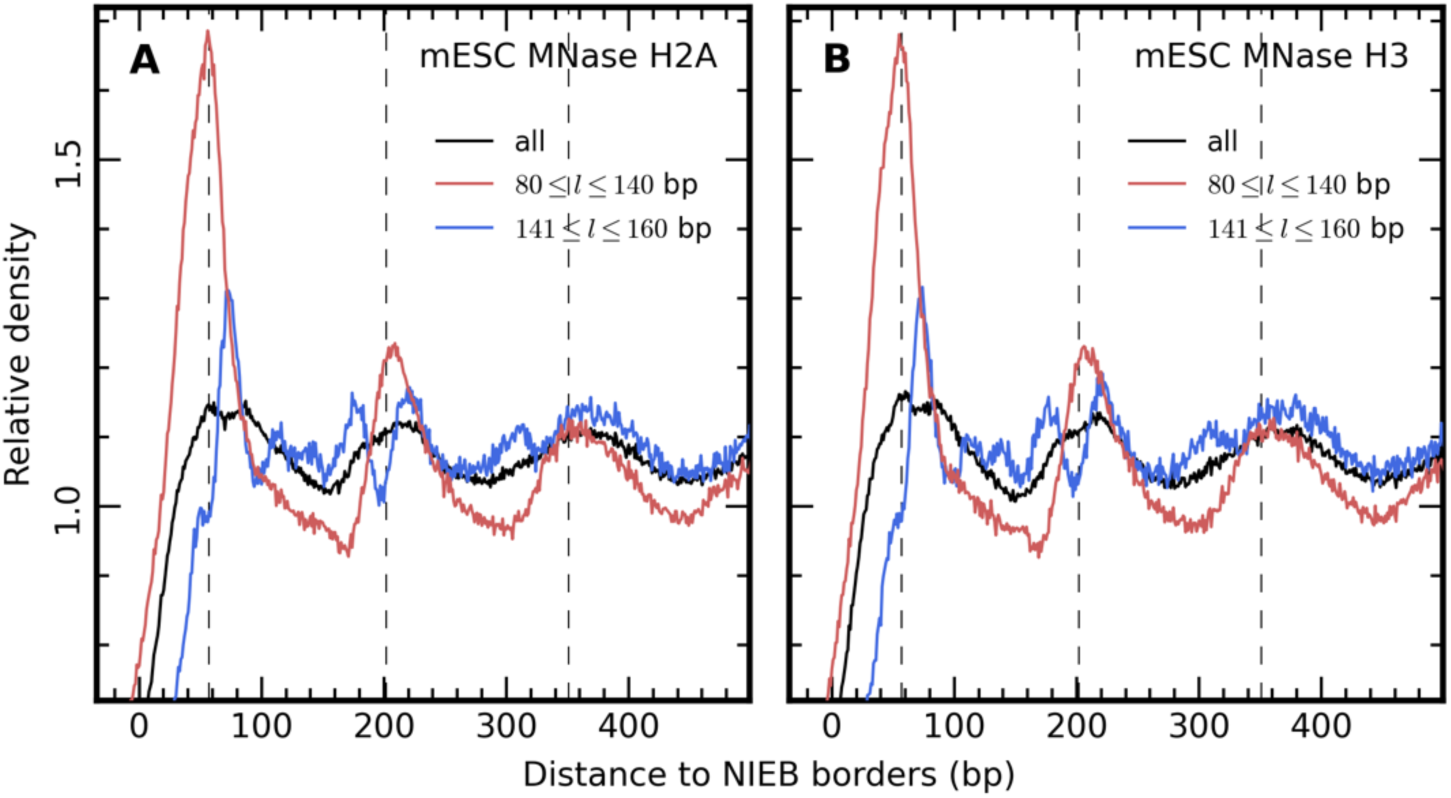
H2A- and H3-associated particles have identical distribution at NIEB borders. A, B Relative MNase ChIP seq density profiles at NIEBs borders for canonical histone H2A (A) and H3 (B) for all (black), subnucleosome-sized (red) and nucleosome-sized (blue) fragments as in Fig 1D.

**Figure EV2.**
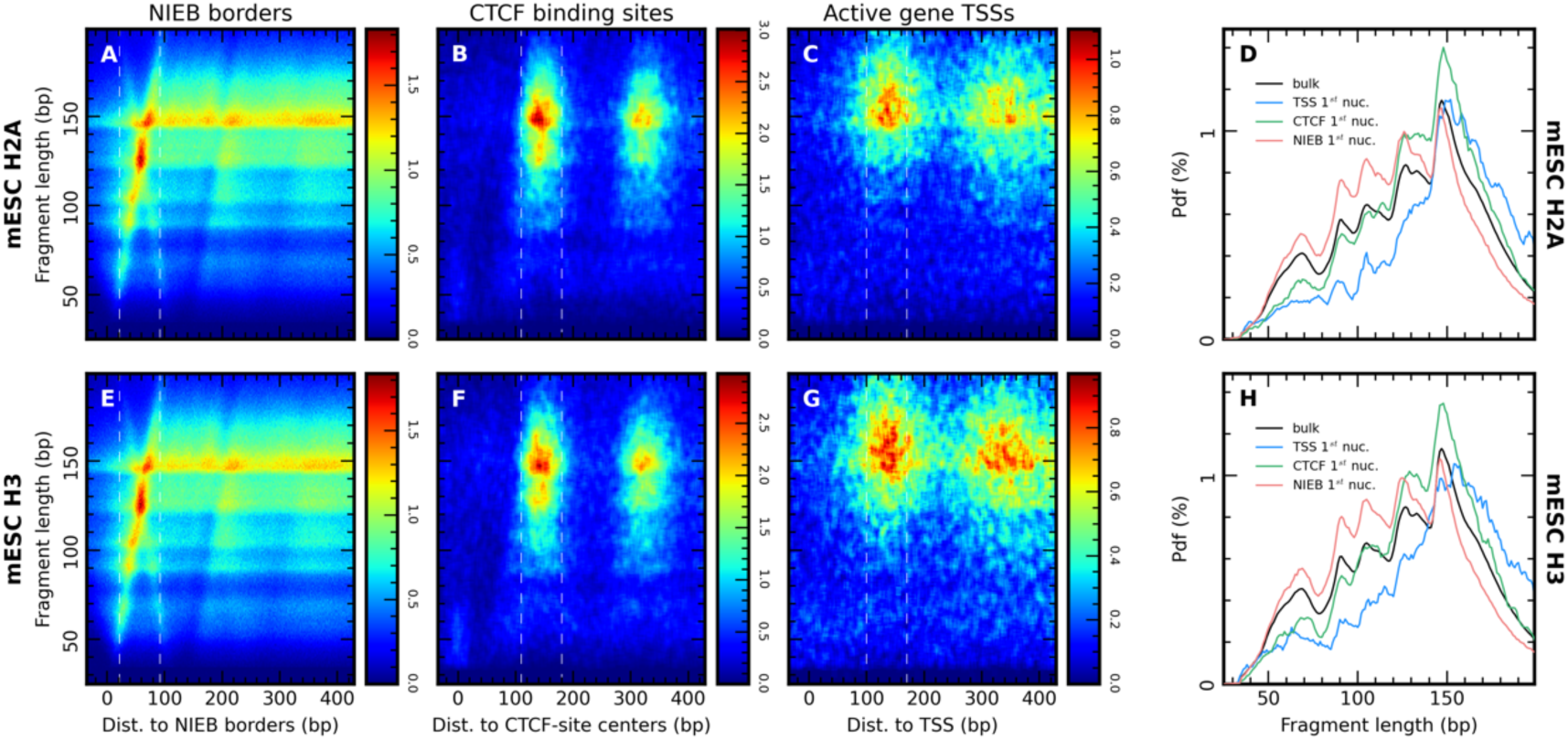
Subnucleosome-sized particles at NIEBs borders are associated to H2A and H3. A-D Same as Fig 2A-D for MNase ChIP seq for canonical histone H2A (mESC H2A data). E-H Same as Fig 2A-D for MNase ChIP seq for canonical histone H3 (mESC H3 data).

**Figure EV3.**
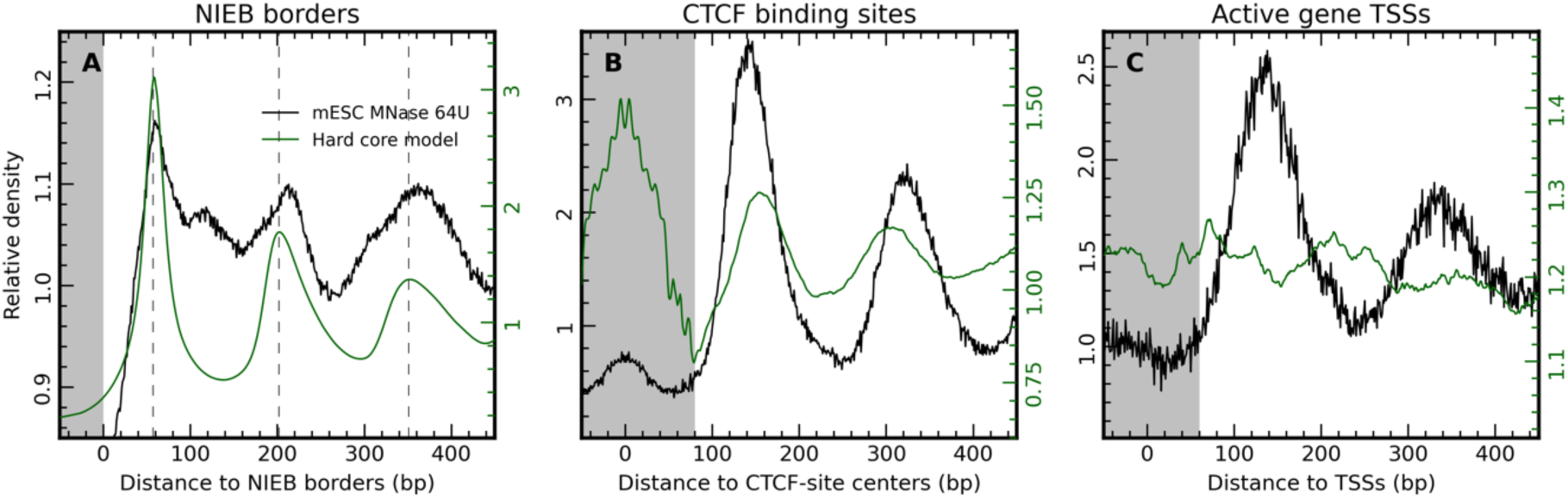
Role of the histone variant H2A.Z in nucleosome fragment length. A-C Relative MNase seq density profiles in mESC (mESC MNase 64U, dotted lines, Y-scale to the left) and predicted nucleosome occupancy profile (full lines, Y-scale to the right) at borders of the NIEBs (A), CTCF binding sites (B) and active gene TSSs (C). The nucleosome depleted region is highlighted in grey. Predicted nucleosome occupancy is calculated from the energy required to form a nucleosome due to DNA bendability and statistical nucleosome positioning. Profiles are computed at 1 bp resolution. Negative (resp. positive) distance values correspond to position inside (resp. outside) the NIEBs.

**Figure EV4.**
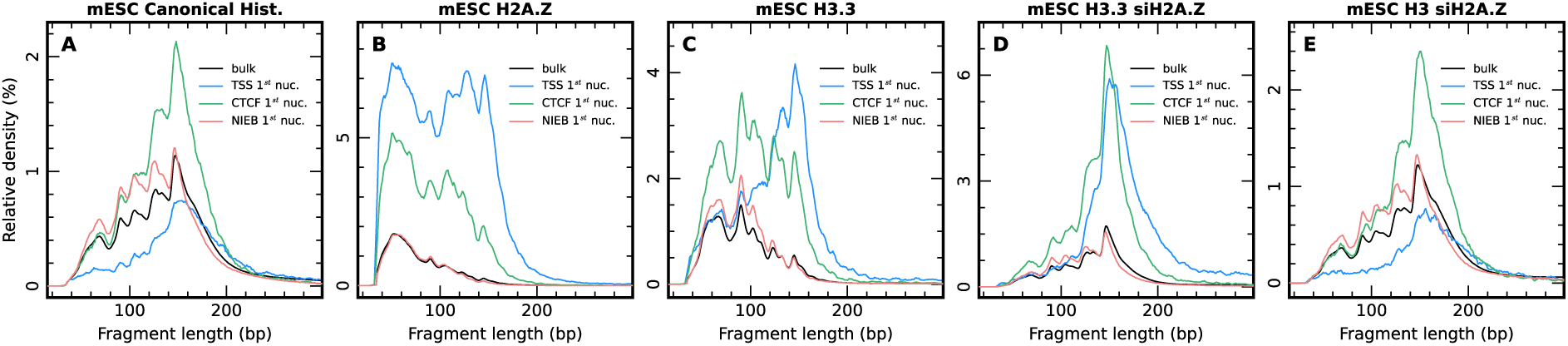
Role of the histone variant H2A.Z in nucleosome fragment length. A-F Relative density as a function of DNA fragment length of MNase ChIP seq for mESC MNase Canonical data (A), mESC H2A.Z data (B), mESC H3.3 data (C), mESC H3.3 siH2A.Z data (D) and mESC H3 siH2A.Z data in the bulk (red) or at NIEBs border (red), CTCF binding site (green) and active TSSs (blue). The fragment size in computed at the position of the 1st nucleosome (highlighted with white dotted lines in Fig 2). It corresponds to the data shown in Fig 2D, H, L, P and T normalized for the genome average fragment density of each experiment. Relative densities are expressed in percent (%).

**Figure EV5.**
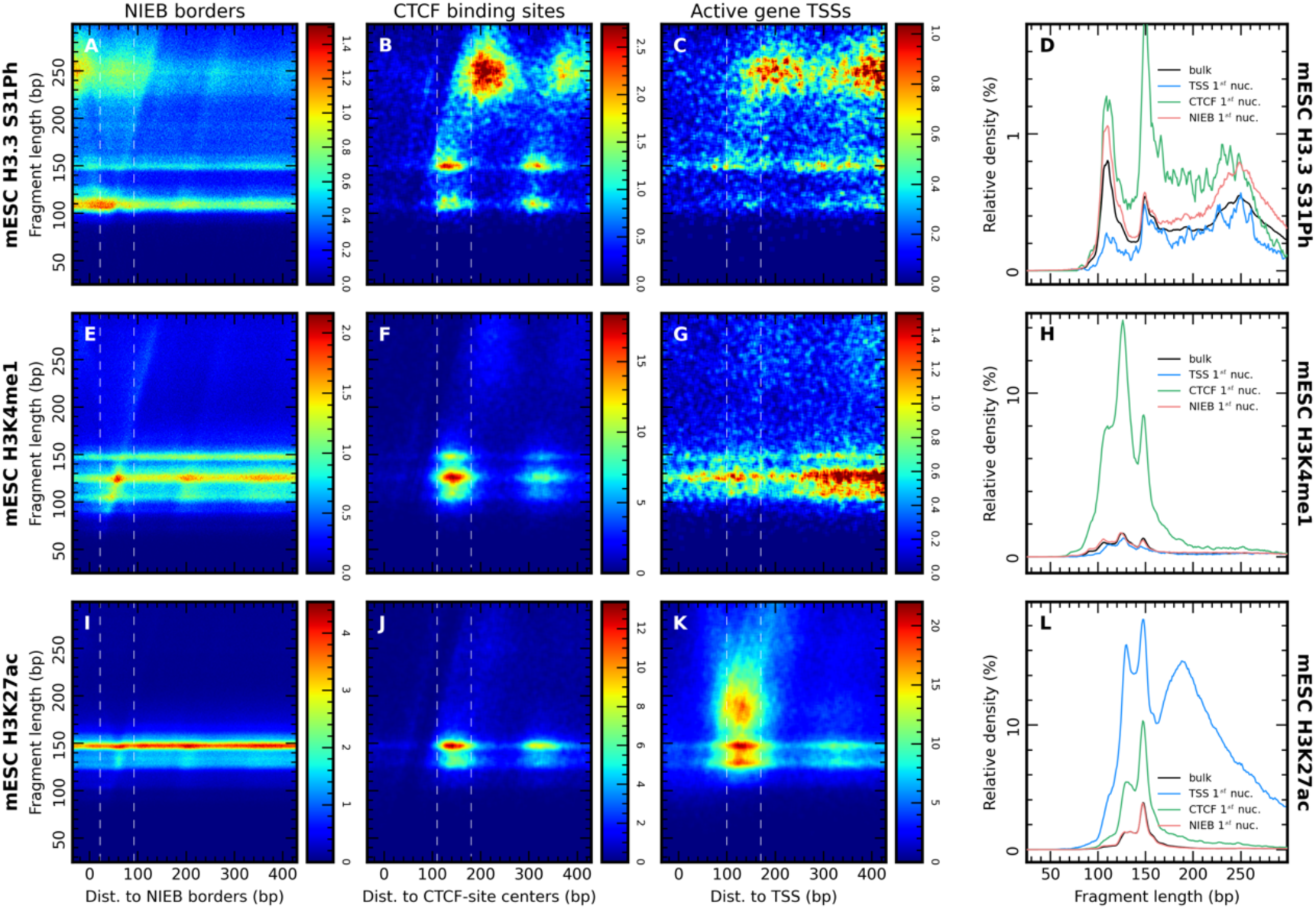
Nucleosome positioning by NIEBs depends on histone tail modifications. A-L Same as Fig 3 but with an enlarged fragment length range to encompass fragments of length 200 bp to 300 bp. D, H, LSame data as in Fig 3D, H and L expressed as fragment relative densities by normalizing for the genome average fragment density of each experiments. Relative densities are expressed in percent (%).

**Figure EV6.**
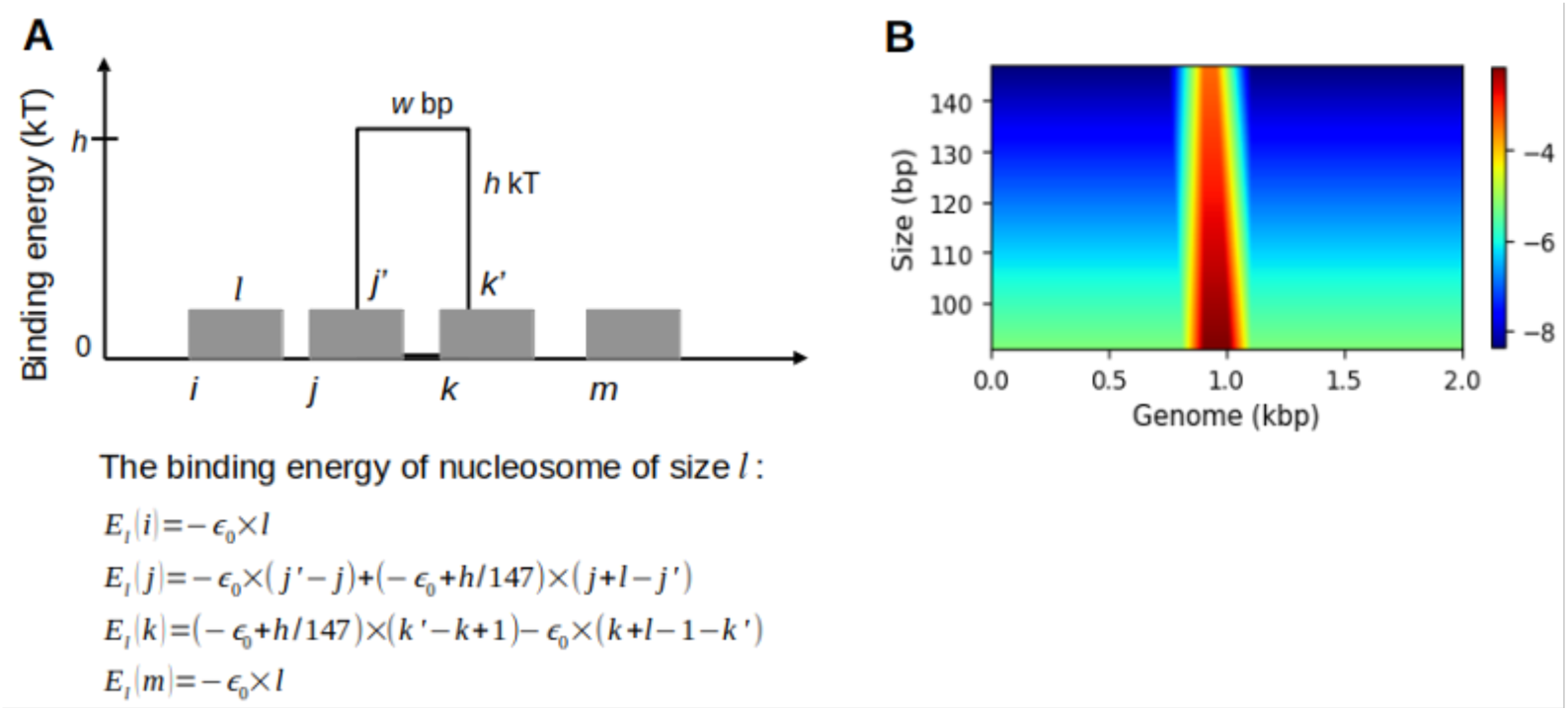
Modeling of partially wrapped nucleosomes at NIEB borders. A A gate-function of width *w* and height h/w is positioned along a flat energy cost per base-pair profile e.i. the only sequence effect is the energy barrier. The energy cost of forming a nucleosome depend on its length = and its overlaping with the gate; four exemplary nucleosomes are shown. B Heatmap representation of the energy matrix *E_l_*(*i*) for the barrier model described in A. Sequence length is *L* = 2 *kb*, NIEB is located between positions 900bp and 1099bp, ∈_0_ = 0.057 kT/bp and ℎ = 5 *kT*.

**Figure EV7.**
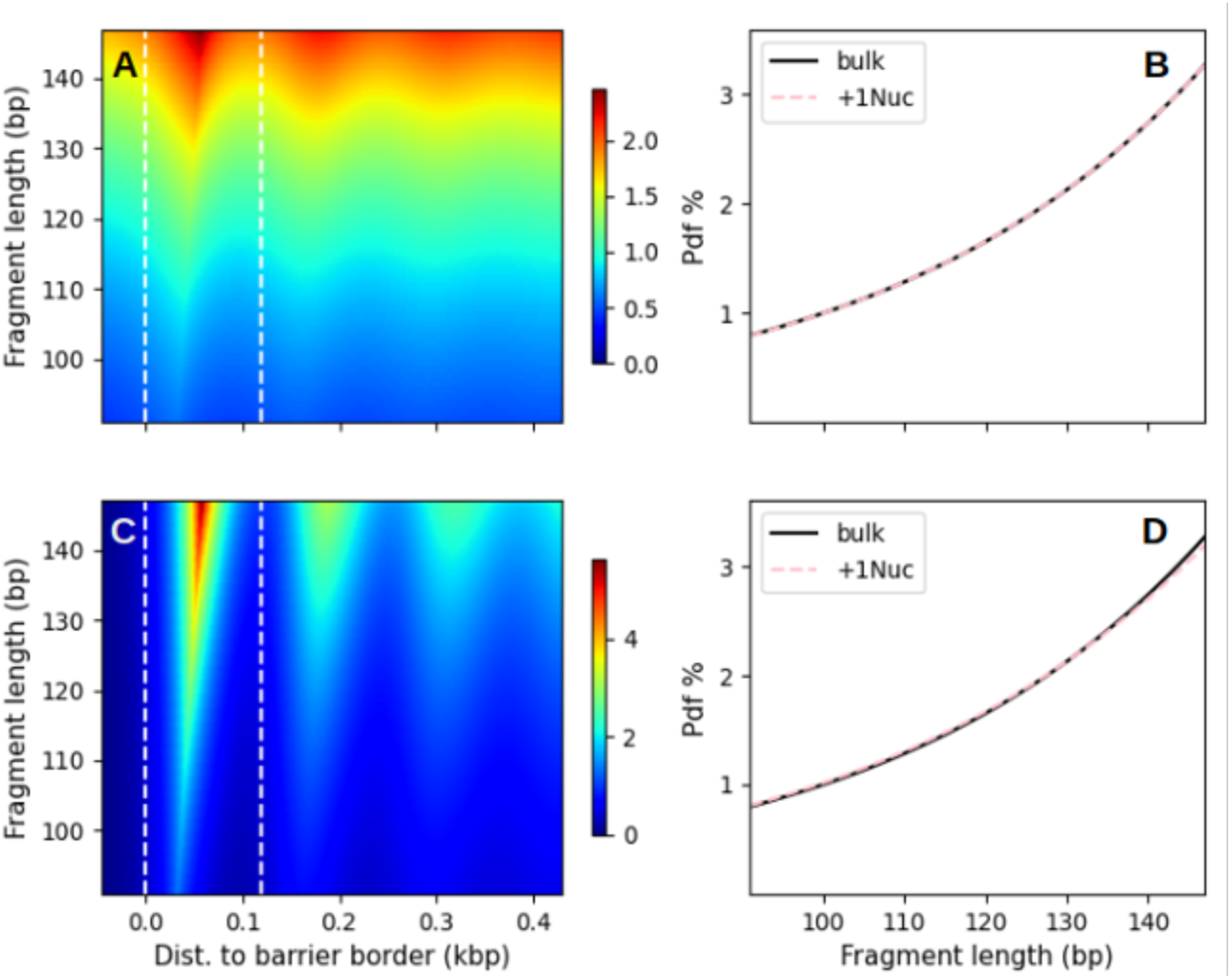
Positioning of partially wrapped nucleosomes at NIEB borders. A, B Same as in Fig 4A, B for a NIEB height ℎ = 2 kT. C, D Same as in Fig 4A, B for a NIEB height ℎ = 10 kT.

## References

Andrew S (2010) Babraham Bioinformatics - FastQC A Quality Control tool for High Throughput Sequence Data.

Armache A, Yang S, Martínez de Paz A, Robbins LE, Durmaz C, Cheong JQ, Ravishankar A, Daman AW, Ahimovic DJ, Klevorn T, et al (2020) Histone H3.3 phosphorylation amplifies stimulation-induced transcription. Nature 583: 852–857

Arneodo A, Drillon G, Argoul F & Audit B (2018) The Role of Nucleosome Positioning in Genome Function and Evolution. In Nuclear Architecture and Dynamics, Lavelle C & Victor J-M (eds) pp 41–79. Boston: Academic Press

Awazu A (2017) Prediction of nucleosome positioning by the incorporation of frequencies and distributions of three different nucleotide segment lengths into a general pseudo k-tuple nucleotide composition. Bioinformatics 33: 42–48

Barbier J, Vaillant C, Volff J-N, Brunet FG & Audit B (2021) Coupling between Sequence-Mediated Nucleosome Organization and Genome Evolution. Genes 12: 851

Barski A, Cuddapah S, Cui K, Roh T-Y, Schones DE, Wang Z, Wei G, Chepelev I & Zhao K (2007) High-Resolution Profiling of Histone Methylations in the Human Genome. Cell 129: 823– 837

Brahma S & Henikoff S (2020) Epigenome Regulation by Dynamic Nucleosome Unwrapping. Trends Biochem Sci 45: 13–26

Brunet FG, Audit B, Drillon G, Argoul F, Volff J-N & Arneodo A (2018) Evidence for DNA Sequence Encoding of an Accessible Nucleosomal Array across Vertebrates. Biophys J 114: 2308–2316

Chereji RV & Morozov AV (2014) Ubiquitous nucleosome crowding in the yeast genome. Proceedings of the National Academy of Sciences 111: 5236–5241

Chevereau G, Arneodo A & Vaillant C (2011) Influence of the genomic sequence on the primary structure of chromatin. Frontiers in Life Science 5: 29–68

Chevereau G, Palmeira L, Thermes C, Arneodo A & Vaillant C (2009) Thermodynamics of Intragenic Nucleosome Ordering. Phys Rev Lett 103: 188103

Clarkson CT, Deeks EA, Samarista R, Mamayusupova H, Zhurkin VB & Teif VB (2019) CTCF-dependent chromatin boundaries formed by asymmetric nucleosome arrays with decreased linker length. Nucleic Acids Res 47: 11181–11196

Drillon G, Audit B, Argoul F & Arneodo A (2015) Ubiquitous human ‘master’ origins of replication are encoded in the DNA sequence via a local enrichment in nucleosome excluding energy barriers. J Phys Condens Matter 27: 064102

Drillon G, Audit B, Argoul F & Arneodo A (2016) Evidence of selection for an accessible nucleosomal array in human. BMC Genomics 17: 526

Eaton ML, Galani K, Kang S, Bell SP & MacAlpine DM (2010) Conserved nucleosome positioning defines replication origins. Genes Dev 24: 748–753

Field Y, Fondufe-Mittendorf Y, Moore IK, Mieczkowski P, Kaplan N, Lubling Y, Lieb JD, Widom J & Segal E (2009) Gene expression divergence in yeast is coupled to evolution of DNA-encoded nucleosome organization. Nat Genet 41: 438–445

He S, Wu Z, Tian Y, Yu Z, Yu J, Wang X, Li J, Liu B & Xu Y (2020) Structure of nucleosome-bound human BAF complex. Science 367: 875–881

van der Heijden T, van Vugt JJFA, Logie C & van Noort J (2012) Sequence-based prediction of single nucleosome positioning and genome-wide nucleosome occupancy. Proceedings of the National Academy of Sciences 109: E2514–E2522

Henikoff JG, Belsky JA, Krassovsky K, MacAlpine DM & Henikoff S (2011) Epigenome characterization at single base-pair resolution. Proc Natl Acad Sci U S A 108: 18318–18323

Isaac RS, Jiang F, Doudna JA, Lim WA, Narlikar GJ & Almeida R (2016) Nucleosome breathing and remodeling constrain CRISPR-Cas9 function. eLife 5: e13450

Jin C, Zang C, Wei G, Cui K, Peng W, Zhao K & Felsenfeld G (2009) H3.3/H2A.Z double variant– containing nucleosomes mark ‘nucleosome-free regions’ of active promoters and other regulatory regions. Nat Genet 41: 941–945

Kaplan N, Moore IK, Fondufe-Mittendorf Y, Gossett AJ, Tillo D, Field Y, LeProust EM, Hughes TR, Lieb JD, Widom J, et al (2009) The DNA-encoded nucleosome organization of a eukaryotic genome. Nature 458: 362–366

Kono H & Ishida H (2020) Nucleosome unwrapping and unstacking. Current Opinion in Structural Biology 64: 119–125

Konrad SF, Vanderlinden W & Lipfert J (2022) Quantifying epigenetic modulation of nucleosome breathing by high-throughput AFM imaging. Biophysical Journal 121: 841–851

Koopmans WJA, Buning R, Schmidt T & van Noort J (2009) spFRET Using Alternating Excitation and FCS Reveals Progressive DNA Unwrapping in Nucleosomes. Biophysical Journal 97: 195–204

Kornberg RD & Stryer L (1988) Statistical distributions of nucleosomes: nonrandom locations by a stochastic mechanism. Nucleic Acids Res 16: 6677–6690

Krietenstein N, Wal M, Watanabe S, Park B, Peterson CL, Pugh BF & Korber P (2016) Genomic Nucleosome Organization Reconstituted with Pure Proteins. Cell 167: 709–721.e12

Kubik S, O’Duibhir E, de Jonge WJ, Mattarocci S, Albert B, Falcone J-L, Bruzzone MJ, Holstege FCP & Shore D (2018) Sequence-Directed Action of RSC Remodeler and General Regulatory Factors Modulates +1 Nucleosome Position to Facilitate Transcription. Mol Cell 71: 89–102.e5

Langmead B & Salzberg SL (2012) Fast gapped-read alignment with Bowtie 2. Nat Methods 9: 357–359

Lee W, Tillo D, Bray N, Morse RH, Davis RW, Hughes TR & Nislow C (2007) A high-resolution atlas of nucleosome occupancy in yeast. Nat Genet 39: 1235–1244

Li S, Wei T & Panchenko AR (2023) Histone variant H2A.Z modulates nucleosome dynamics to promote DNA accessibility. Nat Commun 14: 769

Lobbia VR, Trueba Sanchez MC & van Ingen H (2021) Beyond the Nucleosome: Nucleosome-Protein Interactions and Higher Order Chromatin Structure. Journal of Molecular Biology 433: 166827

Lorch Y, Maier-Davis B & Kornberg RD (2014) Role of DNA sequence in chromatin remodeling and the formation of nucleosome-free regions. Genes Dev 28: 2492–2497

Lowary PT & Widom J (1998) New DNA sequence rules for high affinity binding to histone octamer and sequence-directed nucleosome positioning. J Mol Biol 276: 19–42

Luger K, Mäder AW, Richmond RK, Sargent DF & Richmond TJ (1997) Crystal structure of the nucleosome core particle at 2.8 Å resolution. Nature 389: 251–260

Martin M (2011) Cutadapt removes adapter sequences from high-throughput sequencing reads. EMBnet.journal 17: 10–12

Martire S, Gogate AA, Whitmill A, Tafessu A, Nguyen J, Teng Y-C, Tastemel M & Banaszynski LA (2019) Phosphorylation of histone H3.3 at serine 31 promotes p300 activity and enhancer acetylation. Nat Genet 51: 941–946

Mieczkowski J, Cook A, Bowman SK, Mueller B, Alver BH, Kundu S, Deaton AM, Urban JA, Larschan E, Park PJ, et al (2016) MNase titration reveals differences between nucleosome occupancy and chromatin accessibility. Nat Commun 7: 11485

Milani P, Chevereau G, Vaillant C, Audit B, Haftek-Terreau Z, Marilley M, Bouvet P, Argoul F & Arneodo A (2009) Nucleosome positioning by genomic excluding-energy barriers. Proceedings of the National Academy of Sciences 106: 22257–22262

Polach KJ & Widom J (1995) Mechanism of Protein Access to Specific DNA Sequences in Chromatin: A Dynamic Equilibrium Model for Gene Regulation. Journal of Molecular Biology 254: 130–149

Schwartz U, Németh A, Diermeier S, Exler JH, Hansch S, Maldonado R, Heizinger L, Merkl R & Längst G (2019) Characterizing the nuclease accessibility of DNA in human cells to map higher order structures of chromatin. Nucleic Acids Res 47: 1239–1254

Tillo D & Hughes TR (2009) G+C content dominates intrinsic nucleosome occupancy. BMC Bioinformatics 10: 442

Tompitak M, Barkema GT & Schiessel H (2017a) Benchmarking and refining probability-based models for nucleosome-DNA interaction. BMC Bioinformatics 18: 1–7

Tompitak M, Vaillant C & Schiessel H (2017b) Genomes of Multicellular Organisms Have Evolved to Attract Nucleosomes to Promoter Regions. Biophys J 112: 505–511

Trifonov EN (1985) Curved DNA. CRC Crit Rev Biochem 19: 89–106

Tsankov AM, Thompson DA, Socha A, Regev A & Rando OJ (2010) The role of nucleosome positioning in the evolution of gene regulation. PLoS Biol 8: e1000414

Udugama M, Vinod B, Chan FL, Hii L, Garvie A, Collas P, Kalitsis P, Steer D, Das PP, Tripathi P, et al (2022) Histone H3.3 phosphorylation promotes heterochromatin formation by inhibiting H3K9/K36 histone demethylase. Nucleic Acids Res 50: 4500–4514

Vaillant C, Audit B & Arneodo A (2007) Experiments confirm the influence of genome long-range correlations on nucleosome positioning. Phys Rev Lett 99: 218103

Vaillant C, Palmeira L, Chevereau G, Audit B, d’Aubenton-Carafa Y, Thermes C & Arneodo A (2010) A novel strategy of transcription regulation by intragenic nucleosome ordering. Genome Res 20: 59–67

Valouev A, Johnson SM, Boyd SD, Smith CL, Fire AZ & Sidow A (2011) Determinants of nucleosome organization in primary human cells. Nature 474: 516–520

Wen Z, Zhang L, Ruan H & Li G (2020) Histone variant H2A.Z regulates nucleosome unwrapping and CTCF binding in mouse ES cells. Nucleic Acids Research 48: 5939–5952

Widlund HR, Cao H, Simonsson S, Magnusson E, Simonsson T, Nielsen PE, Kahn JD, Crothers DM & Kubista M (1997) Identification and characterization of genomic nucleosome-positioning sequences. J Mol Biol 267: 807–817

Wong LH, Ren H, Williams E, McGhie J, Ahn S, Sim M, Tam A, Earle E, Anderson MA, Mann J, et al (2009) Histone H3.3 incorporation provides a unique and functionally essential telomeric chromatin in embryonic stem cells. Genome Res 19: 404–414

Yu H, Wang J, Lackford B, Bennett B, Li J & Hu G (2021) INO80 promotes H2A.Z occupancy to regulate cell fate transition in pluripotent stem cells. Nucleic Acids Research 49: 6739–6755

Zhang Y, Moqtaderi Z, Rattner BP, Euskirchen G, Snyder M, Kadonaga JT, Liu XS & Struhl K (2009) Intrinsic histone-DNA interactions are not the major determinant of nucleosome positions in vivo. Nat Struct Mol Biol 16: 847–852

Zhang Z, Wippo CJ, Wal M, Ward E, Korber P & Pugh BF (2011) A packing mechanism for nucleosome organization reconstituted across a eukaryotic genome. Science 332: 977–980

